# Sensitivity of Argentinean isolates of *Leptosphaeria maculans* to Azoxystrobin, Boscalid and Prothioconazole

**DOI:** 10.64898/2026.01.08.698322

**Authors:** Micaela Ester Stieben, Ariadna Aixa Yael Castillo, Fernando Matías Romero, Andres Gárriz, Franco Rubén Rossi

## Abstract

Blackleg, caused by *Leptosphaeria maculans*, is a major disease affecting oilseed rape (*Brassica napus*) worldwide. The use of fungicides is an essential tool for disease management; however, limited information is available regarding *L. maculans* sensitivity to fungicides in Argentina. In this study, the sensitivity of *L. maculans* isolates collected from five oilseed rape-growing regions in Argentina was evaluated against three fungicides with different modes of action: prothioconazole (DMI), azoxystrobin (QoI), and boscalid (SDHI). The effective concentration required to inhibit mycelial growth by 50% (EC_50_) was determined using in vitro assays. Mean EC_50_ values for prothioconazole, azoxystrobin, and boscalid were 0.1545 µg mL^-1^, 0.0349 µg mL^-1^, and 0.1557 µg mL^-1^, respectively. A wider variability in sensitivity was observed for azoxystrobin and boscalid compared to prothioconazole. Additionally, discriminatory dose assays revealed significant differences in growth inhibition among isolates, with some showing reduced sensitivity. Spearman correlation analysis indicated a significant positive association between azoxystrobin and boscalid sensitivity at both low and high doses, suggesting potential cross-resistance. However, no significant correlations were found between prothioconazole and the other fungicides. Furthermore, isolates with reduced sensitivity (RSI < 1) did not exhibit a consistent cross-resistance pattern between QoI and SDHI fungicides, indicating that the observed correlations may be influenced by the overall distribution of sensitive isolates rather than a shared resistance mechanism. These findings provide the first insights into *L. maculans* fungicide sensitivity in Argentina, highlighting the need for continued monitoring to prevent resistance development and ensure effective blackleg management.

## 1 Introduction

Oilseed rape (*Brassica napus*) is a globally important crop, primarily valued for its high-quality oil used in food, biodiesel production, and industrial applications. Its protein-rich meal is also a key animal feed (Correndo et al., 2024). However, this crop is significantly impacted by fungal diseases, especially in regions with high humidity. Among these diseases, blackleg (also termed phoma stem canker disease)-caused by the fungal pathogens *Leptosphaeria maculans* and *L. biglobosa*- is one of the most destructive, affecting a wide range of cruciferous crops (Fitt et al., 2006). Economic losses due to blackleg can range from 5% to 50% of the yield in Australia, Canada and Europe (King et al., 2024). These substantial losses highlight the critical need for effective disease management strategies, where fungicides play a pivotal role. The life cycle of *L. maculans* is complex, involving both sexual and asexual reproduction phases. Ascospores are released during rainy periods in the autumn, coinciding with the sowing season, and are easily dispersed by wind. These spores infect young leaves through stomatal openings, leading to necrotic lesions known as phoma leaf spot (Bousset et al., 2018). The pathogen then grows asymptomatically through the leaf petiole into the stem base, causing blackleg in the spring. This can result in premature lodging and significant yield losses (Huang et al., 2014). While blackleg has been the primary focus due to its impact on yield, recent studies have reported an increase in aerial infections—including pod and seed infections—that may serve as additional inoculum sources, contributing to the disease’s spread (Sprague et al., 2018; Van de Wouw et al., 2016).

In Argentina*, L. maculans* was first reported in 2005 and was initially considered the sole causal agent of blackleg in oilseed rape (*Brassica napus*) (Gaetán, 2005). However, recent studies have identified *L. biglobosa* oilseed rape in fields in Baradero, Buenos Aires (Rossi et al., 2018), indicating the coexistence of both pathogens in the region. This finding has significant implications for disease management strategies. Blackleg is the most prevalent disease in oilseed rape in Argentina, though no studies have yet quantified the economic losses caused by this disease.

Fungicides are a cornerstone in the management of Blackleg worldwide. However, their efficacy is influenced by factors such as chemical formulation, application timing, and the sensitivity of local pathogen populations(Brachaczek et al., 2021; Steed et al., 2007). Regular monitoring of pathogen sensitivity is essential to ensure the continued effectiveness of fungicides (Zhang and Fernando, 2018). A growing global concern is the emergence of resistance to single-site fungicides, including demethylation inhibitors (DMIs) like triazoles, quinone outside inhibitors (QoIs) like strobilurins, and succinate dehydrogenase inhibitors (SDHIs) like carboxamides. These fungicides target specific enzymes—CYP51 for triazoles, cytochrome bc1 for strobilurins, and succinate dehydrogenase for carboxamides—and are widely used in Argentina (FRAC, 2025). The heavy reliance on a limited number of modes of action increases the risk of resistance development in pathogen populations, potentially compromising disease control (Liu, 2014; Zhang and Fernando, 2018). Therefore, monitoring the loss of pathogen sensitivity through the determination of EC_50_ values is a widely used method in laboratory settings, not only for *L. maculans* but also for various other plant pathogens. This approach helps detect shifts in fungicide sensitivity, enabling timely adjustments in management practices to prevent the spread of resistant strains.

In this context, our study aims to evaluate, for the first time in Argentina, the sensitivity of 156 *L. maculans* isolates to three commonly used fungicides: prothioconazole (a DMI), azoxystrobin (a QoI), and boscalid (an SDHI). By assessing the sensitivity profiles of these isolates, we aim to provide valuable insights into the current status of fungicide efficacy and inform more effective disease management strategies. The findings will contribute to optimizing fungicide use in oilseed rape cultivation in Argentina, potentially reducing the risk of resistance development and improving crop yields.

## 2 Materials and Methods

### 2.1 Isolation of Leptosphaeria maculans

Samples were collected in 2020/2021 from oilseed rape plants in production fields, specifically from plants exhibiting symptoms of Phoma leaf spots and blackleg (phoma stem cankers). Sampling was conducted along transects within each field, collecting samples from different points spaced at least 200 meters apart and avoiding field edges. In each geographic region, samples were taken from 5 to 8 fields, and from each field, between 3 and 5 isolates were obtained.The pathogen isolation was performed as described by West et al. (2002). Symptomatic plant tissue segments were placed directly into a moist chamber without prior disinfection and incubated for 2–4 days at 22–24 °C under a 12-hour light/dark cycle. This method promotes the formation of pycnidia, with a distinctive white-pink exudate (cirrus) rich in pycnidiospores, easily observed under a microscope. For each isolation, an inoculation loop was used to transfer spores from the exudate, which were then streaked on potato dextrose agar (PDA) supplemented with antibiotics (streptomycin 15 mg L^-1^, gentamicin 15 mg L^-1^, and tetracycline 12 mg L^-1^). Through this process, 156 *L. maculans* isolates were obtained from five distinct geographic regions in Argentina. It is important to note that at the time of collection, the fungicide use history in these canola fields or other crops was unknown.

### 2.2 Maintenance of Isolates

The obtained isolates were cultured on solid V8 medium (35 mL V8 juice, 700 mg CaCO₃, 7 g agar, 315 mL H₂O) for 10–12 days under the previously described photoperiod and temperature conditions. After full sporulation, 2 x 2 mm plugs were excised from Petri plates and transferred to 1.5 mL microcentrifuge tubes, then covered with either sterile water or 30% glycerol. Isolates preserved in sterile water were used for routine subcultures, while those stored in 30% glycerol at -80°C were reserved for long-term storage.

### 2.3 Molecular characterization

The molecular identification of the 156 isolates was conducted following the protocol described by Liu et al. (2014). Cultures were incubated for 15–20 days, allowing mycelium to fully cover 50 mm Petri plates. Genomic DNA was extracted from the cultures using the EasyPure® Plant Genomic DNA Kit (Transgen Biotech), following the manufacturer’s instructions. Identification of *L. maculans* and *L. biglobosa* pv. brassicae was performed through PCR using a three-primer strategy, which includes a common primer (LmacR 5′-GCAAAATGTGCTGCGCTCCAGG-3′) and a species-specific primer for each pathogen (LmacF 5′-CTTGCCCACCAATTGGATCCCCTA-3′ for *L. maculans* and LbigF 5′-ATCAGGGGATTGGTGTCAGCAGTTGA-3′ for *L. biglobosa*). The PCR cycling conditions were as follows: 95°C for 2 min, followed by 35 cycles of 95°C for 15 s, 68°C for 30 s, and 72°C for 1 min, with a final extension at 72°C for 10 min. This approach yields a 331 bp product for *L. maculans* isolates and a 444 bp product for *L. biglobosa* pv. brassicae isolates. DNA from isolates ME24 of *L. maculans* (kindly provided by Professor Kim Hammond-Kosack, Rothamsted Research-UK) and Tapidor of *L. biglobosa* pv. Brassicae (kindly provided by Professor Bruce Fitt, University of Hertfordshire-UK) were used as positive controls in each assay.

### 2.4 Fungicide Stock Preparation

The fungicides used in all assays were of technical grade (≥95% purity). Stock solutions were prepared and stored at -20°C for both EC_50_ and discriminatory dose determination assays. Prothioconazole-Desthio stocks (hereafter referred to as Prothioconazole) were prepared in acetone at a final concentration of 100 mg mL^-1^, while stocks of Azoxystrobin and Boscalid were prepared in dichloromethane at a final concentration of 200 mg mL^-1^

### 2.5 Determination of Average EC_50_

To assess the sensitivity of the isolates obtained, the effective concentration required to inhibit *L. maculans* growth by 50% (EC_50_) was determined. A total of 40 randomly selected isolates from five Argentine regions were used to establish the average EC_50_ value. The concentration ranges used in EC_50_ determinations for each fungicide were calibrated in a preliminary test, which evaluated maximum inhibitory doses on a subset of selected isolates. Two distinct approaches were applied based on each fungicide’s mode and site of action:

#### Petri Dish Assays

Susceptibility to prothioconazole (DMI) was tested using 150-mm Petri dishes containing potato dextrose agar (PDA) (Van De Wouw et al., 2017). The stock solution was then diluted to achieve final concentrations of 0 (control), 0.01, 0.05, 0.1, 0.5, and 5 μg mL-1. The fungicide solutions were incorporated into molten PDA medium cooled to approximately 50 °C to avoid thermal degradation, and thoroughly mixed before pouring into plates. Isolates were pre-cultured on V8 agar for 10 days at 20 °C in the dark. From each culture, 2 × 2 mm mycelial plugs were taken from the colony margin and transferred to the center of each plate. For each isolate and concentration, two replicate plates were prepared. Plates were incubated at 20 °C in the dark for one week. After incubation, plates were photographed, and colony areas were measured using Image Pro-Plus 4.5 software. Growth inhibition was calculated relative to the control (0 μg mL⁻¹), and EC₅₀ values were estimated from dose-response curves using GraphPad Prism 8 (Siah et al., 2010).

#### Multiwell Plate Assays

To evaluate sensitivity to azoxystrobin and boscalid (QoI and SDHI fungicides), isolates were subcultured as described. Pycnidiospores suspensions were prepared by adding 2 mL of sterile water to each Petri dish, incubating for 2 minutes, and gently scraping the plate surface with a sterile spatula. The resulting homogenate was filtered, centrifuged at 12,000 xg for 10 minutes, and resuspended in 1 mL of sterile water, then stored at -20°C. Prior to inoculation, pycnidiospore concentrations were adjusted to 5 x 10^5^ spore mL^-1^ using a counting chamber. EC_50_ determinations were performed in 96-well microtiter plates, with each well containing 50 µL of clarified V8 medium (Fortune et al., 2024) with streptomycin (50 µg mL^-1^) and final fungicide concentrations of 0 (Control), 0.0001, 0.001, 0.01, 0.1, and 1, , , µg mL^-1^. A 50 µL spore suspension was added to each well for a final concentration of 2.5 x 10^5^ spore mL^-1^. Each isolate was tested in quadruplicate at each fungicide concentration. Plates were incubated at 20°C in darkness without shaking for 5 days. Spore germination and growth were measured by absorbance at 405 nm using a BioTek Synergy microplate reader. EC_50_ values for each isolate were determined through growth inhibition analysis from dose-response curves using GraphPad Prism 8 (Siah et al., 2010).

### 2.6 Fungicide Sensitivity of Isolates

To evaluate the sensitivity of 156 isolates to prothioconazole, azoxystrobin, and boscalid, two discriminatory doses were selected based on the average EC_50_ values obtained from the 40 previously tested isolates. As with EC_50_ determinations, two methodologies were used based on the fungicide mode of action and target site:

#### Petri Dish Assays

Sensitivity to prothioconazole (DMI) was tested by inoculating fresh cultures on 150 mm Petri dishes with 8-9 isolates per plate, following previous cultivation protocols. A discriminatory dose test was conducted on PDA medium supplemented with prothioconazole at two concentrations (0.1 and 0.5 μg mL^-1^). Each isolate and concentration were tested in duplicate, with plates incubated for one week under previously described conditions. Fungal growth areas were measured using Image Pro-Plus 4.5 software, and the inhibition of growth was calculated as a percentage of mycelial area reduction compared to the control:

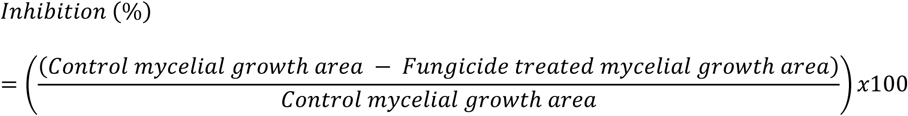

#### Multiwell Plate Assays

For azoxystrobin and boscalid (QoI and SDHI), sensitivity was evaluated using fresh cultures in 96-well plates, similar to the previous protocol. Two discriminatory doses (0.1 and 1 µg mL^-1)^ were used for both fungicides. Growth inhibition in the presence of fungicides was determined by averaging absorbance readings from four technical replicates for each condition. Growth inhibition was calculated as a percentage of absorbance reduction compared to the control:

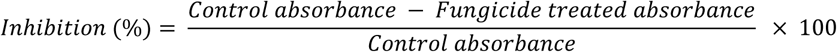

### 2.7 Relative Sensitivity Index (RSI) Calculation

To assess the relative sensitivity of each isolate within the tested population, we calculated a Relative Sensitivity Index (RSI). For each fungicide and concentration tested, RSI was defined as the ratio between the growth inhibition (%) of a given isolate and the mean growth inhibition (%) observed for the entire population at that same concentration. This index expresses the response of each isolate in relation to the population average under identical conditions.

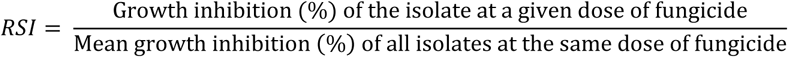

RSI values greater than 1 indicate isolates with higher-than-average sensitivity, while values below 1 reflect lower sensitivity relative to the population. This approach was chosen due to the absence of a known wild-type reference isolate in Argentina and the lack of historical isolates predating the use of the active ingredients evaluated in this study. The RSI metric allows for a comparative analysis of isolate behavior without assuming the availability of a sensitive baseline strain.

## 3 Results

### 3.1 Isolation and Molecular Characterization

A collection of 156 isolates of the blackleg causal agent was obtained from leaf and stem samples displaying typical disease symptoms, collected from oilseed rape fields across five geographical regions in Argentina. Four of these regions are located in the province of Buenos Aires, while the fifth corresponds to the province of Entre Ríos. These areas were selected due to their long-standing history of oilseed rape cultivation (Figure 1). Molecular identification confirmed that all isolates belonged to *L. maculans*, as evidenced by the amplification of a single PCR product of 331 bp, specific to this species (data not shown).

**Figure 1.**
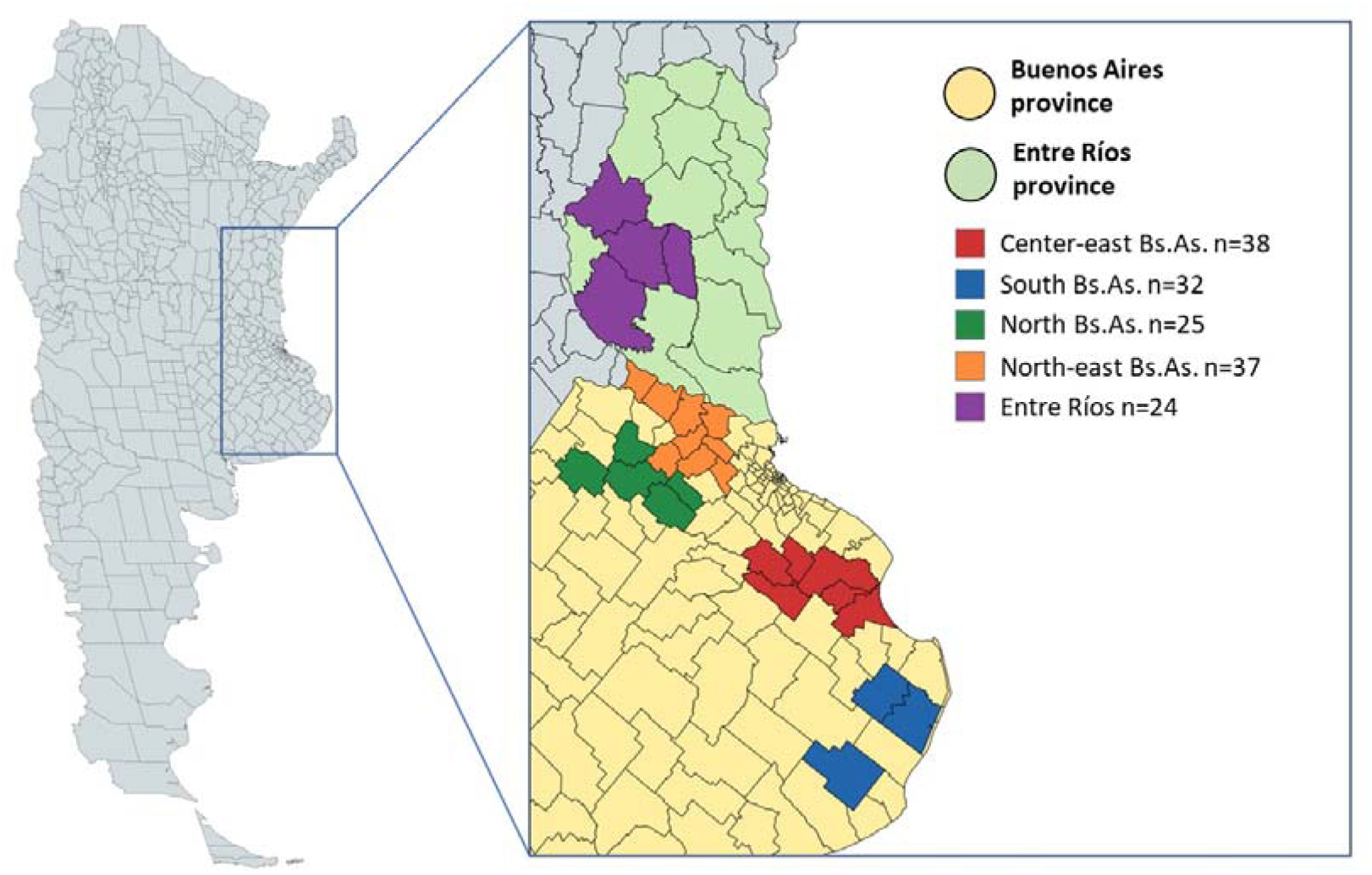
Map of Buenos Aires and Entre Ríos province (Argentina) showing the geographical origin of *L. maculans* isolates collected between 2021 and 2023. The different colours represent the five sampling regions: Center-east Buenos Aires (CE, Bs.As.), North-east Buenos Aires (NE, Bs.As.), North Buenos Aires (N, Bs.As.), South Buenos Aires (S, Bs.As.), and Entre Ríos (ER).

Of the 156 isolates collected (Figure 1), 116 were tested against all three fungicides. However, 133 isolates were evaluated for sensitivity to prothioconazole, while 142 isolates were assessed for azoxystrobin and 140 for boscalid. This variation in the number of isolates tested for each fungicide was due to experimental constraints and availability of isolates in sufficient quantity for all assays.

### 3.2 Determination of EC_50_

To date, no studies assessing fungicide sensitivity in isolates of the blackleg causal agent have been conducted in Argentina or South America. Given this lack of information, we determined the EC_50_ of 40 randomly selected isolates representing five major oilseed rape-growing regions in Argentina. To achieve this, three technical-grade fungicides with distinct modes and sites of action were tested to estimate the EC_50_ for each isolate. The EC_50_ values for individual isolates ranged from 0.0012 to 3.666 µg mL⁻¹ for prothioconazole, 0.0003 to 0.720032 µg mL⁻¹ for azoxystrobin, and 0.0007 to 1.5157 µg mL⁻¹ for boscalid. Based on the individual EC_50_ values of the 40 isolates analyzed, the mean EC_50_ was 0.1545 µg mL⁻¹ for prothioconazole, 0.0349 µg mL⁻¹ for azoxystrobin, and 0.1557 µg mL⁻¹ for boscalid (Table 1). Regionally, isolates from north Buenos Aires exhibited the highest average EC_50_ values for prothioconazole and boscalid, whereas isolates from the central-east region of Buenos Aires showed the highest mean EC_50_ value for azoxystrobin (Table 1).

**Table 1.** EC_50_ range for 40 *L. maculans* isolates from five regions in Argentina (2021–2023) evaluated against fungicides (prothioconazole, azoxystrobin, and boscalid). The table presents the EC_50_ range and the mean EC_50_ values for each fungicide. *N* represents the number of isolates analyzed.

| Region | *N* | EC<sub>50</sub> (µg mL<sup>-1</sup>) |  |  | Mean EC<sub>50</sub> (µg mL<sup>-1</sup>) |  |  |
| --- | --- | --- | --- | --- | --- | --- | --- |
|  |  | Azoxystrobin | Boscalid | Prothioconazole | Azoxystrobin | Boscalid | Prothioconazole |
| *CE Bs.As.* | 10 | 0.0004 –<br>0.7200 | 0.0025 –<br>0.9770 | 0.0012 – 0.2410 | 0.0841 | 0.1769 | 0.0732 |
| *ER* | 9 | 0.0027 –<br>0.0700 | 0.0099 –<br>0.2700 | 0.0114 – 0.1790 | 0.0242 | 0.1200 | 0.0649 |
| *NE Bs.As.* | 10 | 0.0024 –<br>0.0980 | 0.0270 –<br>0.6630 | 0.0046 – 0.1140 | 0.0242 | 0.1480 | 0.0477 |
| *S Bs.As.* | 6 | 0.0003 –<br>0.0170 | 0.0100 –<br>0.1010 | 0.0025 – 0.1180 | 0.0073 | 0.0564 | 0.0534 |
| *N Bs.As.* | 5 | 0.0020 –<br>0.0320 | 0.0007 –<br>1.5157 | 0.0529 – 3.6660 | 0.0105 | 0.3110 | 0.7919 |
| **Total** | 40 | 0.0003 –<br>0.7200 | 0.0007 –<br>1.5157 | 0.0012 – 3.6660 | 0.0349 | 0.1557 | 0.1545 |

An analysis of isolate distribution across EC_50_ ranges revealed that 19 isolates (48.7%) of the 40 analyzed isolates had an EC_50_ between 0.051 and 0.2 µg mL^-1^ for prothioconazole (Figure 2A). For azoxystrobin, 24 isolates (60%) exhibited EC_50_ values within the 0.0051–0.05 µg mL⁻¹ range (Figure 2B). Finally, 21 isolates (52.5%) had EC_50_ values between 0.0301 and 0.2 µg mL⁻¹ for boscalid (Figure 2C).

**Figure 2.**
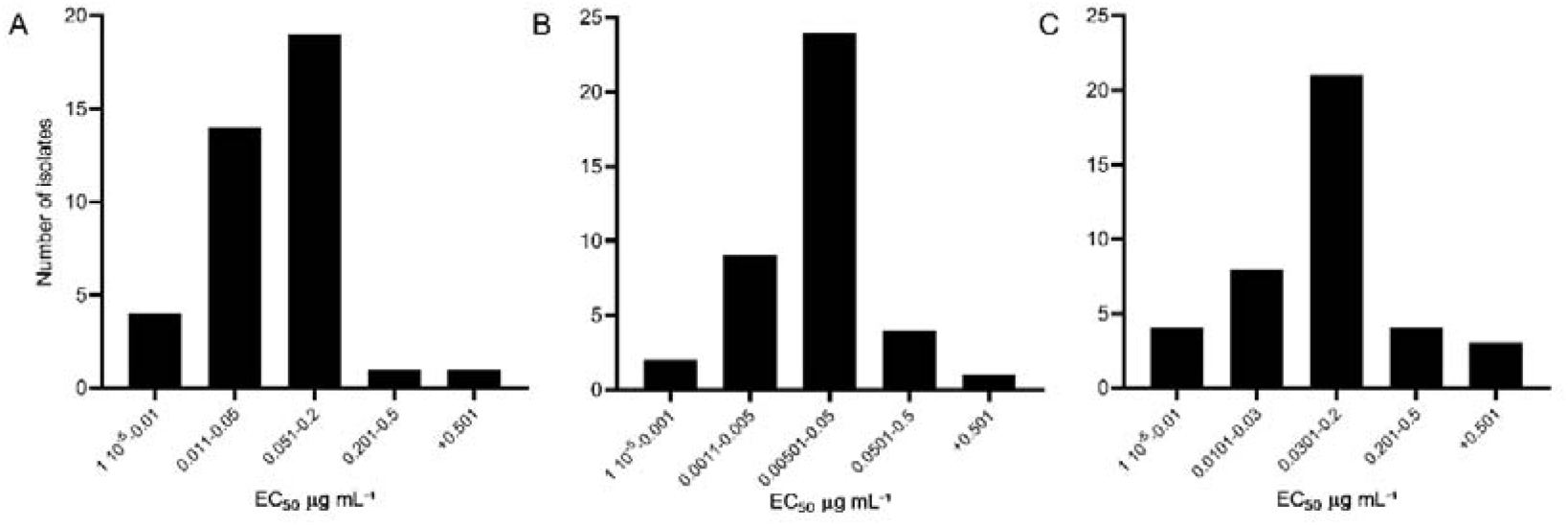
Frequency distribution of the effective concentration (EC₅₀) of three fungicides in 40 isolates of *L. maculans* collected from different geographical regions in Argentina. EC₅₀ values were determined through *in vitro* assays, for the fungicide prothioconazole (A), mycelial growth inhibition was evaluated using an PDA medium assay. The inhibition of pycnidiospore germination and mycelial growth, was measured photometrically at 405 nm for the fungicides azoxystrobin (B) and boscalid (C).

### 3.3 Sensitivity of isolates to prothioconazole

Based on the observed EC_50_ values for each fungicide, we conducted an exploratory analysis to assess the sensitivity of a larger number of isolates (n = 133) collected from the same regions previously evaluated. For this purpose, a discriminatory dose assay was performed for each fungicide, analyzing the percentage of growth inhibition at two concentrations per fungicide. The sensitivity to prothioconazole was assessed by supplementing PDA agar medium with 0.1 µg mL^-1^ and 0.5 µg mL^-1^ of the fungicide. At 0.1 µg mL^-1^, growth inhibition ranged from 0.06% to 96.73%, with an average inhibition of 65.71%. At 0.5 µg mL^-1^, inhibition values increased, ranging from 5.22% to 100%, with a mean inhibition of 89.49%. The comparative analysis between these concentrations revealed a significant increase in fungicidal effectiveness at 0.5 µg mL^-1^ (Figure 3A). The distribution of inhibition percentages at 0.1 µg mL^-1^ showed a slight negative skew (−0.66), while at 0.5 µg mL^-1^, the skewness was more pronounced (−3.53), accompanied by high kurtosis (18.41), suggesting a greater concentration of isolates exhibiting high inhibition. These results indicate a clear shift in sensitivity with increasing prothioconazole concentration, as most isolates at 0.5 µg mL^-1^ displayed high growth inhibition, particularly accumulating in the 90–100% inhibition range (Figure 3B and C).

**Figure 3.**
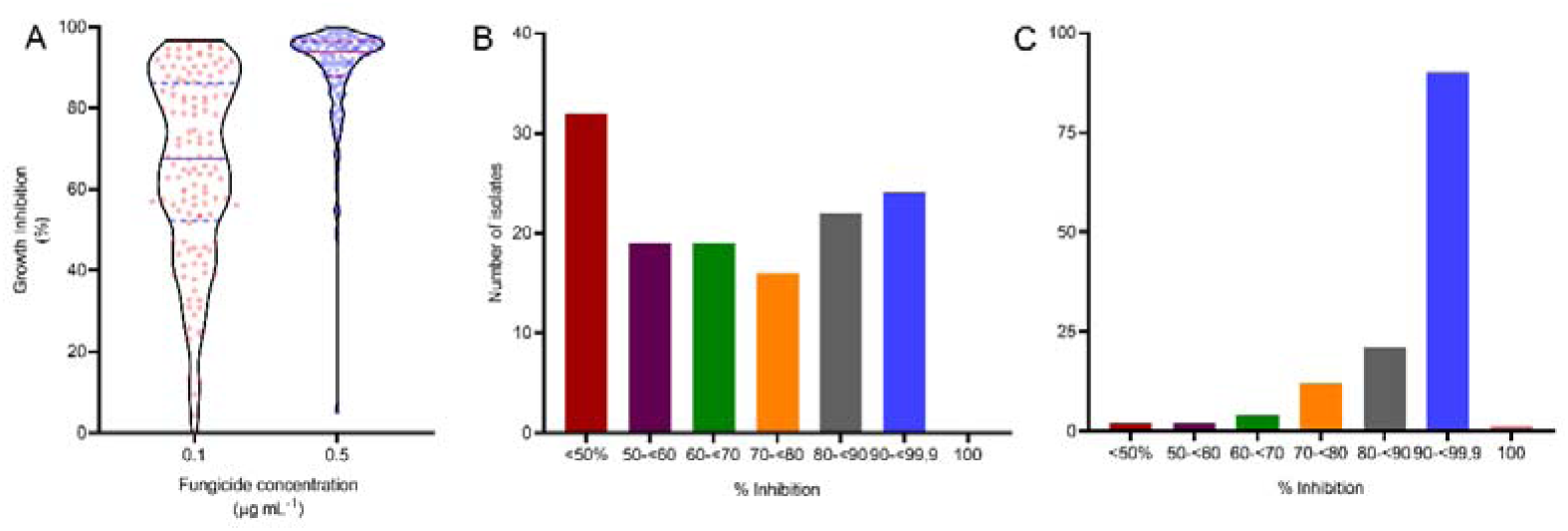
Effect of prothioconazole on the mycelial growth of 133 *L. maculans* isolates. The distribution of mycelial growth inhibition in response to two discriminatory doses of 0.1 and 0.5 µg mL^-1^ in PDA medium is shown. Growth inhibition is expressed as a percentage relative to a control treatment without prothioconazole. The violin plot in panel (A) represents the distribution of inhibition across isolates for each discriminatory dose tested (0.1 and 0.5 µg mL^-1^). Results are presented as means ± standard error (SE) from 133 isolates. A Shapiro-Wilks normality test indicated that data did not follow a normal distribution, so a non-parametric analysis was performed using the Mann-Whitney test, which showed a significant difference (****, P ≤ 0.0001). Additionally, the number of isolates falling within different inhibition ranges for the two discriminatory doses of prothioconazole was determined: 0.1 µg mL^-1^ (B) and 0.5 µg mL^-1^ (C).

Statistical analysis confirmed a significant difference in inhibition distribution between the two concentrations, reinforcing the dose-dependent response of isolates to prothioconazole (Figure 3A). Furthermore, the regional analysis of growth inhibition revealed that isolates from the northeastern region of Buenos Aires exhibited greater sensitivity to prothioconazole, showing higher inhibition levels at both 0.1 µg mL^-1^ and 0.5 µg mL^-1^ compared to isolates from other regions (Supplementary Figure 1).

### 3.4 Sensitivity of isolates to Azoxystrobin

The sensitivity to azoxystrobin was evaluated by assessing spore germination in microplates containing clarified V8 medium supplemented with two discriminatory doses of 0.1 µg mL^-1^ and 1 µg mL^-1^ of the active ingredient. At 0.1 µg mL^-1^, growth inhibition ranged from 0% to 74.4%, with an average inhibition of 37.21%. When the concentration was increased to 1 µg mL^-1^, inhibition values ranged from 2.19% to 86.61%, with an overall mean inhibition of 52.28% across all isolates (Figure 4A). A comparative analysis between these concentrations revealed a significant increase in efficacy at the higher dose (Figure 4A). The inhibition data at 0.1 µg mL^-1^ showed an approximately symmetric distribution (skewness = -0.18), whereas at 1 µg mL^-1^, the data exhibited a stronger negative skew (−1.06) and higher kurtosis (1.44), indicating a concentration of values toward higher inhibition percentages. This suggests a dose-dependent response, as most isolates at 1 µg mL^-1^ displayed greater inhibition, with a notable accumulation in the 50–90% inhibition range (Figure 4B and C). Furthermore, the analysis of growth inhibition based on the geographic origin of the isolates showed no significant differences in sensitivity to azoxystrobin. This effect was consistent across both tested concentrations, 0.1 µg mL^1^ and 1 µg mL^-1^ (Supplementary Figure 2).

**Figure 4.**
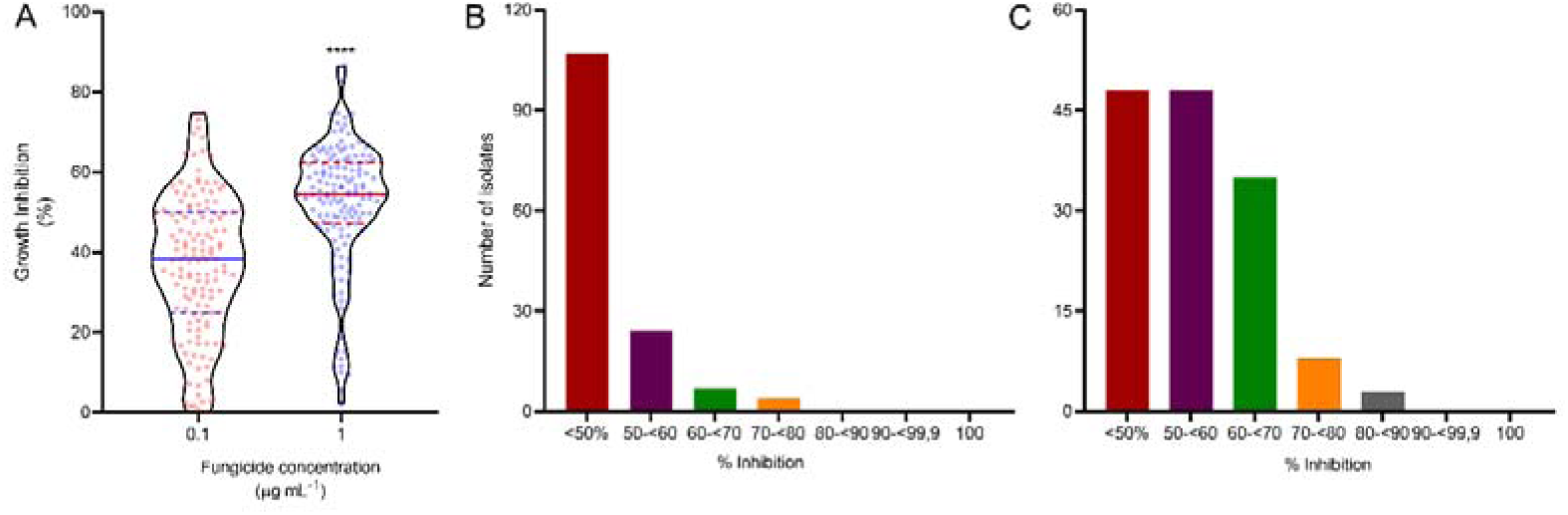
Effect of azoxystrobin on the percentage of mycelial growth inhibition in 142 *L. maculans* isolates. The inhibition of pycnidiospore germination and mycelial growth, measured photometrically at 405 nm, is shown in response to two discriminatory doses of 0.1 and 1 µg mL^-1^ of azoxystrobin in liquid V8 medium using a microtitration assay. Growth inhibition is expressed as a percentage relative to a control treatment without azoxystrobin. The violin plot in panel (A) represents the distribution of inhibition across isolates for each discriminatory dose tested (0.1 and 1 µg mL^-1^). Results are presented as means from 142 isolates. A Shapiro-Wilks normality test indicated that data did not follow a normal distribution, so a non-parametric analysis was performed using the Mann-Whitney test, which showed a significant difference (****, P ≤ 0.0001). Additionally, the number of isolates falling within different inhibition ranges for the two discriminatory doses of azoxystrobin was determined: 0.1 µg mL^-1^ (B) and 1 µg mL^-1^ (C).

### 3.5 Sensitivity of isolates to Boscalid

The sensitivity of isolates to boscalid was evaluated following the same methodology described for azoxystrobin, with culture media supplemented with 0.1µg mL^-1^ and 1 µg mL^-1^ of the active ingredient. The analysis of fungal growth inhibition revealed a clear dose-dependent response, with significantly higher effectiveness observed at the elevated concentration (Figure 5A). At 0.1 µg mL^-1^, the data exhibited slight positive skewness (0.52), indicating a higher frequency of isolates with lower inhibition levels. In contrast, at 1 µg mL^-1^, the skewness shifted to a minor negative value (−0.38), suggesting a more balanced distribution of inhibition across isolates. The addition of 0.1 µg mL^-1^ of boscalid resulted in growth inhibition ranging from 0% to 87.62%, with an overall average inhibition of 30.98%. At 1 µg mL^-1^, the range expanded to 0%–94.89%, with a mean inhibition of 52.52% (Figure 5A). The broader inhibition range at 1 µg mL^-1^ highlights the increased growth suppression of isolates at the higher dose. Statistical analysis (P ≤ 0.0001) further supports the enhanced efficacy of boscalid at elevated concentrations, underlining its potential as an effective fungicide (Figure 5B, C). When analyzing the sensitivity of isolates by geographical origin, no significant differences were observed at either concentration of boscalid (0.1 µg mL^-1^ or 1 µg mL^-1^), suggesting uniform sensitivity across regions (Supplementary Figure 3)

**Figure 5.**
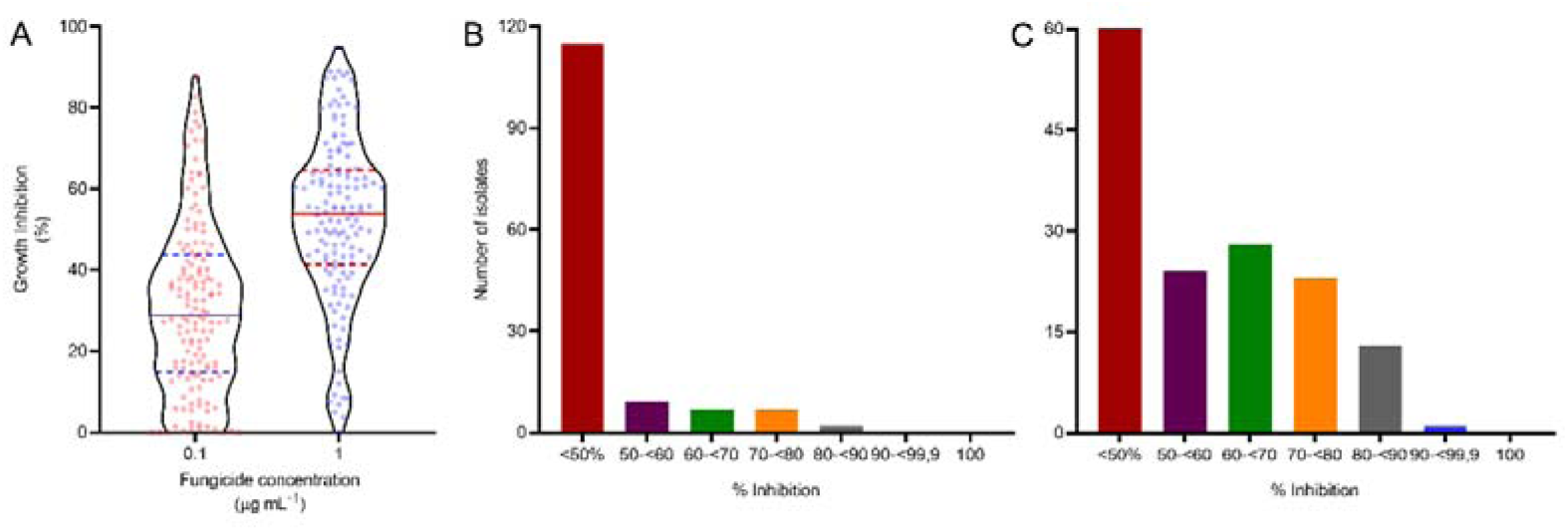
Effect of boscalid on spore germination and mycelial growth of 140 *L. maculans* isolates. The inhibition of pycnidiospore germination and mycelial growth, measured photometrically at 405 nm, can be observed in response to the inclusion of two discriminatory doses (0.1 and 1 µg mL^-1^) of boscalid in liquid V8 medium in a microtitration assay. Growth inhibition is expressed as a percentage relative to a control treatment without boscalid. The violin plot in panel (A) indicates the abundance of samples regarding the inhibition of isolates within each of the discriminatory doses used (0.1 and 1 µg mL^-1^). The results correspond to the means of 140 isolates. A normality test was performed using the Shapiro-Wilks test, which showed that the data did not follow a normal distribution. Therefore, a non-parametric analysis was conducted between both discriminatory doses using the Mann-Whitney test, which revealed a significant difference of ****, P ≤ 0.0001. Additionally, the number of isolates within different inhibition ranges for the two discriminatory doses of boscalid tested was determined: 0.1 µg mL^-1^ (B) and 1 µg mL^-1^ (C).

### 3.6 Resistance Factor Analysis and Cross-Sensitivity Among Fungicides

Relative Sensitivity Index (RSI) analysis revealed variability in fungicide sensitivity among the isolates tested (Table 2). Most isolates exhibited RSI values greater than 1, indicating higher-than-average sensitivity to the fungicides. In contrast, a subset of isolates showed RSI values lower than 1, suggesting reduced sensitivity relative to the population mean.Among the fungicides evaluated, boscalid showed the highest proportion of isolates with RSI < 1 at both tested concentrations (64.31% at 0.1 µg mL^-1^ and 46.55% at 1 µg mL^-1^), followed by azoxystrobin (45.68% at 0.1 µg mL^-1^ and 37.93% at 1 µg mL^-1^) and prothioconazole, which had the lowest proportion of isolates with RSI < 1 (46.55% at 0.1µg mL^-1^ and 29.31% at 0.5 µg mL^-1^). When analyzed at higher concentrations, the proportion of isolates with RSI < 1 decreased, indicating that higher fungicide doses were more effective at inhibiting fungal growth across most isolates.

**Table 2.** Relative Sensibity Index (RSI) analysis in fungicide sensitivity among the isolates tested. The table shows the range of RSI, number and percentage of isolates with RSI ≤1 and ≥1 for each fungicide dose assayed

| Fungicide | Range of<br>RSI | Number of isolates<br>RSI>1 | % | Number of isolates<br>RSI<1 | % |
| --- | --- | --- | --- | --- | --- |
| *Prothioconazole 0.1 µg mL<sup>-1</sup>* | 0,00 –1,50 | 62 | 53,44% | 54 | 46,55% |
| *Prothioconazole 0.5 µg mL<sup>-1</sup>* | 0,06 –1,12 | 82 | 70,68% | 34 | 29,31% |
| ***Azoxystrobin 0.1 µg mL<sup>-1</sup>*** | 0,00 –1,99 | 63 | 54,31% | 53 | 45,68% |
| ***Azoxystrobin 1 µg mL<sup>-1</sup>*** | 0, 04–1,64 | 72 | 62,06% | 44 | 37,93% |
| ***Boscalid 0.1 µg mL<sup>-1</sup>*** | 0,00 –2,94 | 53 | 45,68% | 63 | 64.31% |
| ***Boscalid 1 µg mL<sup>-1</sup>*** | 0,00 –1,88 | 62 | 53,44% | 54 | 46,55% |

Spearman correlation analysis revealed distinct relationships between fungicide sensitivities at both low and high doses. At low concentrations, a weak but significant positive correlation was observed between azoxystrobin and boscalid (r=0.3071, p=0.0008), whereas no significant correlations were found between azoxystrobin and prothioconazole (r=0.0063, p=0.9469) or boscalid and prothioconazole (r=−0.0935, p=0.3180) (Figure 6A-C). At higher fungicide concentrations, the correlation between azoxystrobin and boscalid increased to a moderate level (r=0.4648, p<0.0001), while no significant associations were observed between azoxystrobin and prothioconazole (r=0.1403, p=0.1330) or boscalid and prothioconazole (r=0.1468, p=0.1158) (Figure 6D-F). These results indicate that the response of *L. maculans* isolates to prothioconazole is independent of their sensitivity to azoxystrobin and boscalid, whereas azoxystrobin and boscalid share a more consistent relationship, particularly at higher concentrations.

**Figure 6.**
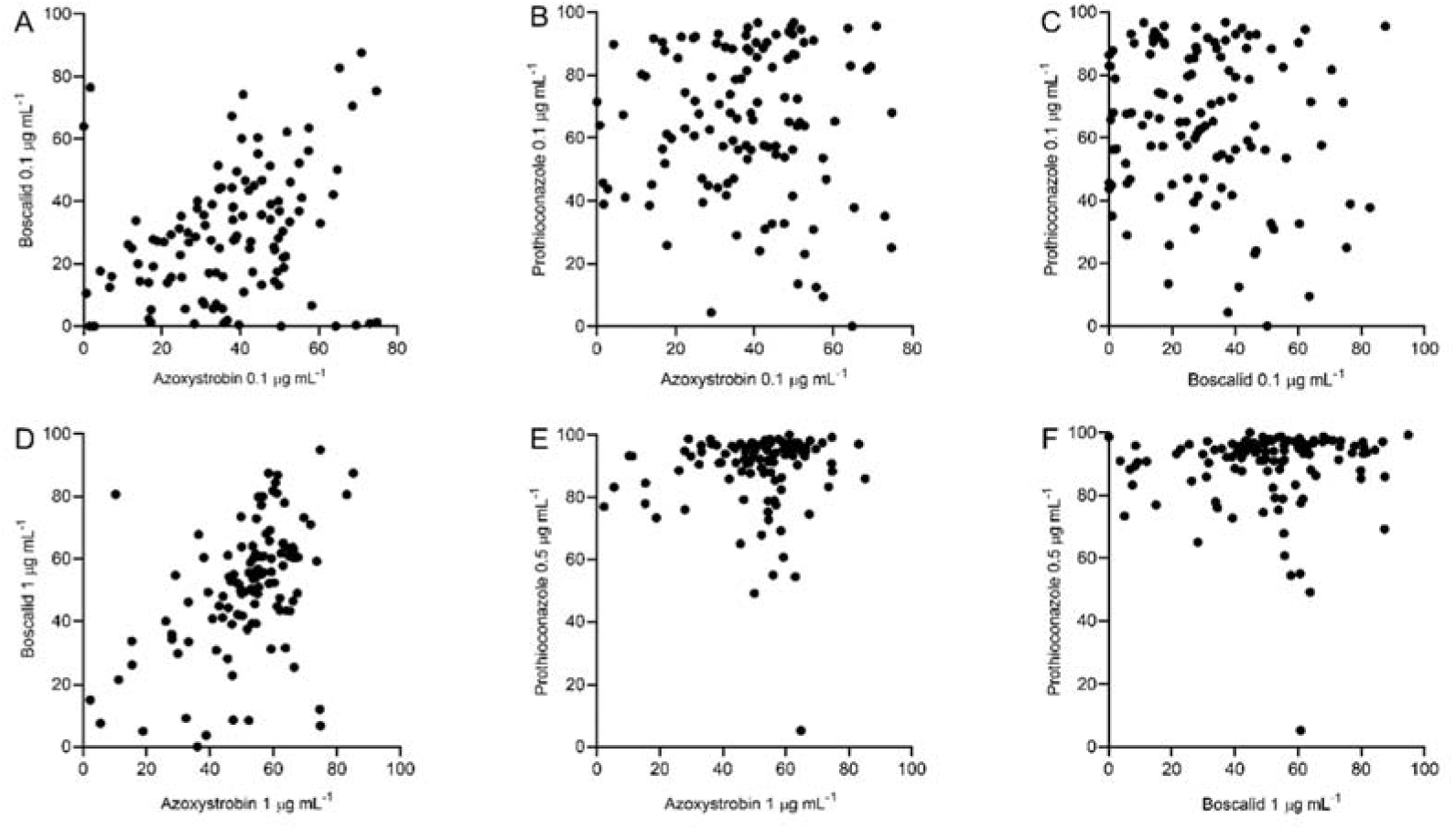
Cross-resistance analysis of *L. maculans* isolates to different fungicides. (A–C) Mycelial growth inhibition at low discriminatory doses (0.1 µg mL^-1^ for azoxystrobin, boscalid, and prothioconazole) in 116 isolates collected from five geographical regions of Argentina. (D–F) Mycelial growth inhibition at high discriminatory doses (1 µg mL^-1^ for azoxystrobin and boscalid, 0.5 µg mL^-1^ for prothioconazole).

However, further analysis considering only isolates with RSI < 1 (indicative of reduced sensitivity) provided a more nuanced interpretation. When evaluating the subset of isolates with RSI < 1, no significant correlations were detected between prothioconazole and azoxystrobin at both low (27 isolates, p>0.05) and high concentrations (13 isolates, p>0.05), nor between prothioconazole and boscalid at low (27 isolates, p>0.05) and high concentrations (14 isolates, p>0.05). This suggests that resistance to prothioconazole occurs independently of the sensitivity levels to the other two fungicides. Interestingly, among isolates with RSI < 1, a significant correlation was observed between azoxystrobin and boscalid at low concentrations (39 isolates, p<0.05), but this correlation was not maintained at high concentrations (Supplementary Table 1).

## 4 Discussion

This study represents the first assessment of fungicide sensitivity in *Leptosphaeria maculans* isolates from Argentina, providing essential data on the efficacy of three key fungicides—prothioconazole, azoxystrobin, and boscalid—used for blackleg disease management. Our results reveal significant variability in sensitivity among isolates and geographic regions, highlighting the importance of local monitoring to guide effective disease control strategies. These findings contribute to a growing global dataset on *L. maculans* fungicide sensitivity and provide a foundation for improved management strategies in Argentina, where limited data on fungicide efficacy against blackleg are available. Until this study, the only scientific evidence on blackleg control in oilseed rape in Argentina reported that fungicides based on azoxystrobin (Amistar®, Syngenta) and propiconazole (Tilt®, Syngenta), applied at the 2–4 leaf stage, significantly reduced stem canker severity (Edwards Molina et al., 2017). Additionally, the only fungicide currently registered in Argentina for blackleg control is Amistar Top® (azoxystrobin + difenoconazole, Syngenta) (CASAFE, 2025). Although oilseed rape cultivation has a history in Argentina and is often adopted as a winter crop alternative to wheat, its planted area has fluctuated considerably over time, according to official data reported by FAO (2025). However, certain regions have consistently grown oilseed rape (Figure 1), and blackleg disease has been reported in these areas (Gaetán, 2005; Rossi et al., 2018; Van de Wouw et al., 2023). Despite this, limited information exists on the susceptibility of different oilseed rape hybrids and varieties to the disease in Argentina. Furthermore, the lack of awareness among producers regarding blackleg management has been associated with significant yield losses (personal observations and communications with oilseed rape producers, 2020–2023). Given that oilseed rape is not a priority crop in Argentina, the origin of *L. maculans* inoculum remains uncertain. However, it is likely that the pathogen persists in weedy *Brassicaceae* species closely related to oilseed rape, such as *Brassica napus* (wild type), *Raphanus sativus*, *Rapistrum rugosum*, *Hirschfeldia incana*, and *Brassica rapa*, which may serve as primary inoculum sources (Chen and Séguin-Swartz, 1999; Oreja et al., 2024). In this context, the present study provides critical information that can directly impact blackleg disease management through fungicide applications and potentially support the registration of new active ingredients currently not approved for use in Argentina.

The variability in fungicide sensitivity observed in *L. maculans* isolates from Argentina highlights the complexity of blackleg management in the region. Among the fungicides tested, prothioconazole exhibited the lowest variability in EC_50_ values, whereas azoxystrobin and boscalid showed a broader distribution, indicating a more heterogeneous response within the population (Figure 2, Table 1 and 2). This pattern aligns with findings from other major oilseed rape-producing regions, such as Australia, Canada, and Europe, where *L. maculans* populations have demonstrated varying degrees of sensitivity to different fungicides (Fajemisin et al., 2022a; Sewell et al., 2017; A. P. Van De Wouw et al., 2021; Wang et al., 2020). When compared to international data, the mean EC_50_ for prothioconazole (0.1545 µg mL^-1^) was within the range reported in previous studies, reinforcing the continued efficacy of DMI fungicides against *L. maculans* (Fajemisin et al., 2022a; Sewell et al., 2017). However, the presence of isolates with elevated EC_50_ values, particularly in northern Argentina, suggests early indications of reduced sensitivity in certain subpopulations. This phenomenon has been documented in other fungal pathogens where prolonged and repeated applications of DMI fungicides have led to gradual shifts in sensitivity (Cools and Fraaije, 2013; A. P. Van De Wouw et al., 2021; Van De Wouw et al., 2017). Given the importance of DMIs in blackleg control, monitoring these shifts is critical to ensure their continued effectiveness.

The response of *L. maculans* isolates to discriminatory fungicide doses further supports these observations (Figures 3, 4, and 5). While high in contrast, azoxystrobin and boscalid exhibited greater variability in sensitivity among isolates, with some showing reduced inhibition compared to global averages. The mean EC_50_ for azoxystrobin in our study (0.0349 µg mL^-1^) was lower than the EC_50_ values reported for pyraclostrobin, another QoI fungicide, in *L. maculans* isolates from Canada (Fraser et al., 2017). This suggests that Argentine isolates remain relatively sensitive to QoIs, though the broader EC_50_ range indicates that selection pressure could lead to reduced sensitivity in the future. Conversely, the mean EC_50_ for boscalid (0.1557 µg mL^-1^) was significantly higher than values reported for European populations (Fajemisin et al., 2022b), suggesting that some Argentine isolates might have reduced sensitivity to SDHI fungicides. Similar trends were observed when comparing EC_50_ values for azoxystrobin with dimoxystrobin (Fajemisin et al., 2022b), reinforcing the possibility that QoI resistance mechanisms could be emerging in certain isolates.

Fungicide concentrations generally resulted in strong inhibition across most isolates, the variability at lower doses suggests that sensitivity to azoxystrobin and boscalid is more variable than to prothioconazole. The violin plots (Figures 3A, 4A, 5A) clearly illustrate this distribution, with a higher concentration of isolates in the lower inhibition ranges for azoxystrobin and boscalid, while prothioconazole displayed a more uniform inhibition response across isolates. This trend aligns with reports indicating that QoI and SDHI fungicides often exhibit greater variability in effectiveness due to differences in resistance mechanisms, including target site mutations and metabolic degradation pathways (Fernández-Ortuño et al., 2017; Lucas et al., 2015). Additionally, the classification of isolates into discrete inhibition categories (Figures 3B–C, 4B–C, 5B–C) provided further insights into the population’s response to each fungicide. At low concentrations, a substantial proportion of isolates exhibited inhibition levels below 50% for azoxystrobin and boscalid, whereas at high concentrations, most isolates showed inhibition levels above 70%, albeit with a wider distribution for these two fungicides. This suggests that while higher doses effectively control most isolates, the broader distribution at lower doses may indicate the early stages of reduced sensitivity in some subpopulations.

A significant finding of this study was the presence of regional differences in fungicide sensitivity. Isolates from the northeastern region of Buenos Aires exhibited significantly higher sensitivity to prothioconazole compared to those from other regions (Figure 3 and Supplementary Figure 1). Similar reports have been documented in Argentina for other fungal pathogens, such as *Zymoseptoria tritici* and *Ramularia collo-cygni* (Augusti et al., 2020; Erreguerena et al., 2022). Even for *L. maculans*, regional differences in DMI fungicide sensitivity have been observed in other parts of the world (Fajemisin et al., 2022a; A. P. Van De Wouw et al., 2021; Van De Wouw et al., 2017).

While the central objective of this study was not to determine the mechanisms underlying potential fungicide resistance but rather to explore population-level sensitivity, it is likely that genetic factors contribute to the observed variations in DMI fungicide sensitivity (Price et al., 2015). These genetic modifications, which drive variations in sensitivity, may also be influenced by external factors related to agronomic practices. First, historical fungicide application practices vary across Argentina, with some regions potentially experiencing higher selection pressure due to the repeated use of the same active ingredients. Second, environmental conditions such as temperature and humidity can influence fungal population dynamics and fungicide effectiveness. Lastly, differences in agronomic practices, including crop rotation, cultivar selection, and disease management strategies, may contribute to region-specific variability in sensitivity (Angela P Van De Wouw et al., 2021; West et al., 2001). Notably, no significant regional differences were observed for azoxystrobin or boscalid sensitivity, suggesting that selection pressures affecting DMI fungicides may not similarly impact QoI or SDHI fungicides. This could be due to differences in the mode of action of these fungicides, the frequency of their use, or the underlying genetic resistance mechanisms specific to *L. maculans* populations in Argentina. Further research, including molecular characterization of fungicide resistance markers, is needed to better understand the factors shaping regional sensitivity patterns.

Spearman correlation analysis revealed a significant positive correlation between azoxystrobin and boscalid sensitivity at both low and high doses, suggesting a potential link in their resistance mechanisms. Azoxystrobin belongs to the quinone outside inhibitors (QoIs), which target the cytochrome bc1 complex (complex III) in the mitochondrial electron transport chain, leading to the disruption of ATP synthesis. Similarly, boscalid is a succinate dehydrogenase inhibitor (SDHI) that acts on complex II of the mitochondrial electron transport chain, impairing fungal respiration. The shared role of these fungicides in disrupting mitochondrial respiration may explain the observed cross-sensitivity. In our study, the mean EC_50_ values (Table 1) for azoxystrobin (0.0349 µg mL^-1^) and boscalid (0.1557 µg mL^-1^) displayed high variability among isolates, with some showing reduced sensitivity to both fungicides. At low fungicide concentrations (0.1 µg mL^-1^), a weak but significant positive correlation was observed between azoxystrobin and boscalid sensitivity (r = 0.3071, p = 0.0008), while at higher concentrations (1 µg mL^-1^), this correlation increased to a moderate level (r = 0.4648, p < 0.0001) (Figure 6). However, studies evaluating cross-sensitivity patterns in *L. maculans* populations from other regions have reported contrasting findings. For example, recent data from the Czech Republic did not show a positive correlation between QoI and SDHI sensitivity (Fajemisin et al., 2022b), although dimoxystrobin was evaluated instead of azoxystrobin in that study. Similarly, studies on *Botrytis cinerea* and *Ascochyta rabiei* populations found no significant correlation between QoI and SDHI fungicides (Kim and Xiao, 2010; Wise et al., 2008), whereas a positive correlation has been reported in *Zymoseptoria tritici* populations (Lavrukaitė et al., 2024). A more detailed evaluation of our data (Table 2), focusing on isolates with reduced sensitivity (RSI < 1), suggests that the observed correlation may not necessarily indicate a shared resistance mechanism but rather an effect driven by the overall distribution of sensitivity levels across the population. When analyzing only isolates with RSI < 1 for both azoxystrobin and boscalid, the correlation between these fungicides disappeared at higher concentrations (Supplementary Table 2). This finding implies that the positive correlation observed in the full dataset may be influenced by the predominance of highly sensitive isolates rather than reflecting a direct association between resistance mechanisms. These results are consistent with the significant increase in growth inhibition across all isolates as azoxystrobin and boscalid concentrations increased (Figures 4A and 5A). In other words, while general population trends suggest an association between QoI and SDHI sensitivity, resistant isolates do not necessarily exhibit cross-resistance. These findings emphasize the need to distinguish between overall population trends and resistance-specific mechanisms when interpreting fungicide sensitivity data. In contrast, no significant correlations were found between prothioconazole and the other fungicides (Figure 6), indicating that DMI fungicides may still be an effective alternative for managing isolates with reduced sensitivity to QoI or SDHI compounds, and vice versa. The analysis of RSI further supports this observation, as the proportion of isolates with RSI < 1 for prothioconazole was considerably lower than for azoxystrobin and boscalid (Table 2). This suggests that prothioconazole remains a reliable option for controlling *L. maculans* populations in Argentina. Additionally, correlation analysis among isolates with RSI < 1 revealed no significant associations between prothioconazole and azoxystrobin or between prothioconazole and boscalid at either fungicide concentration (Supplementary Table 1). This lack of correlation indicates that resistance mechanisms affecting QoI and SDHI fungicides do not necessarily confer reduced sensitivity to DMI fungicides, reinforcing their potential as alternative treatment options. These findings align with previous reports where resistance to DMIs has been shown to develop independently of resistance to QoI or SDHI fungicides in other fungal pathogens (Kim and Xiao, 2010; Wise et al., 2008).

Unfortunately, the limited availability of data on the sensitivity or resistance of *L. maculans* to the fungicides evaluated in this study (prothioconazole, azoxystrobin, and boscalid) complicates decision-making regarding their potential use in commercial fungicide formulations. However, in general terms, and with a few exceptions, most isolates exhibited sensitivity to one, two, or even all three fungicides tested. This suggests that blackleg disease could be effectively managed through the application of fungicide mixtures combining DMI + QoI, DMI + SDHI, or even DMI + QoI + SDHI. Fungicide resistance management strategies recommend using fungicides with different modes of action rather than relying on a single fungicide class to control plant diseases. Given the limited data available on fungicide efficacy in Argentina, our findings provide a valuable reference for the development of blackleg management strategies.

Overall, our findings indicate that *L. maculans* populations in Argentina remain largely sensitive to prothioconazole, azoxystrobin, and boscalid, with only a few isolates exhibiting reduced sensitivity. This suggests that, at present, these fungicides can still be used effectively for blackleg management. The data presented in this study provide a valuable baseline for understanding the current fungicide sensitivity landscape in Argentina, a country where blackleg research and management strategies remain underdeveloped compared to other major oilseed rape-producing regions. However, the detection of isolates with reduced sensitivity highlights the need for further investigation into the underlying mechanisms of resistance and the potential selection pressures driving these changes. Future studies should focus on molecular characterization of resistance mechanisms, in-field assessments of fungicide efficacy, and evaluations of genetic susceptibility to *L. maculans* in commercially available oilseed rape hybrids. Additionally, long-term monitoring programs will be essential to track potential shifts in fungicide sensitivity over time and to ensure the continued effectiveness of chemical control strategies. Integrating these efforts with improved agronomic practices and host resistance management will be key to developing a sustainable approach for blackleg control in Argentina.

## 5 Acknowledgements

FMR, AG, and FRR are members of the Research Career of CONICET, Argentina. SME and AAYC are doctoral fellow of FONCyT, Argentina. The authors are grateful to Beatriz Wiss (CPA-CONICET, Argentina) and Juan Ezquiaga (CIC, Buenos Aires province, Argentina) for their technical support and assistance during field sampling and laboratory analyses. We are also grateful to the farmers who allowed access to their fields for sample collection.

## 6 Funding

This work was financially supported by grants of Agencia Nacional de Promoción Científica y Tecnológica (PICT Startup 2020-0053 and PICT 2019-1137) and CONICET (PIBAA 28720210100989CO) to FRR.

## 7 Author Contributions

M.E.S. and A.A.Y.C. performed sample collection, conducted laboratory assays, and contributed to manuscript writing. F.M.R. and A.G. participated in data interpretation and manuscript drafting. F.R.R. conceived and led the research project, secured funding, and wrote the manuscript. All authors reviewed and approved the final version of the manuscript.

## Notes

### Competing Interest Statement

The authors have declared no competing interest.

