## Supplemental Figure 1 for "Sensitivity of Argentinean isolates of *Leptosphaeria maculans* to Azoxystrobin, Boscalid and Prothioconazole"

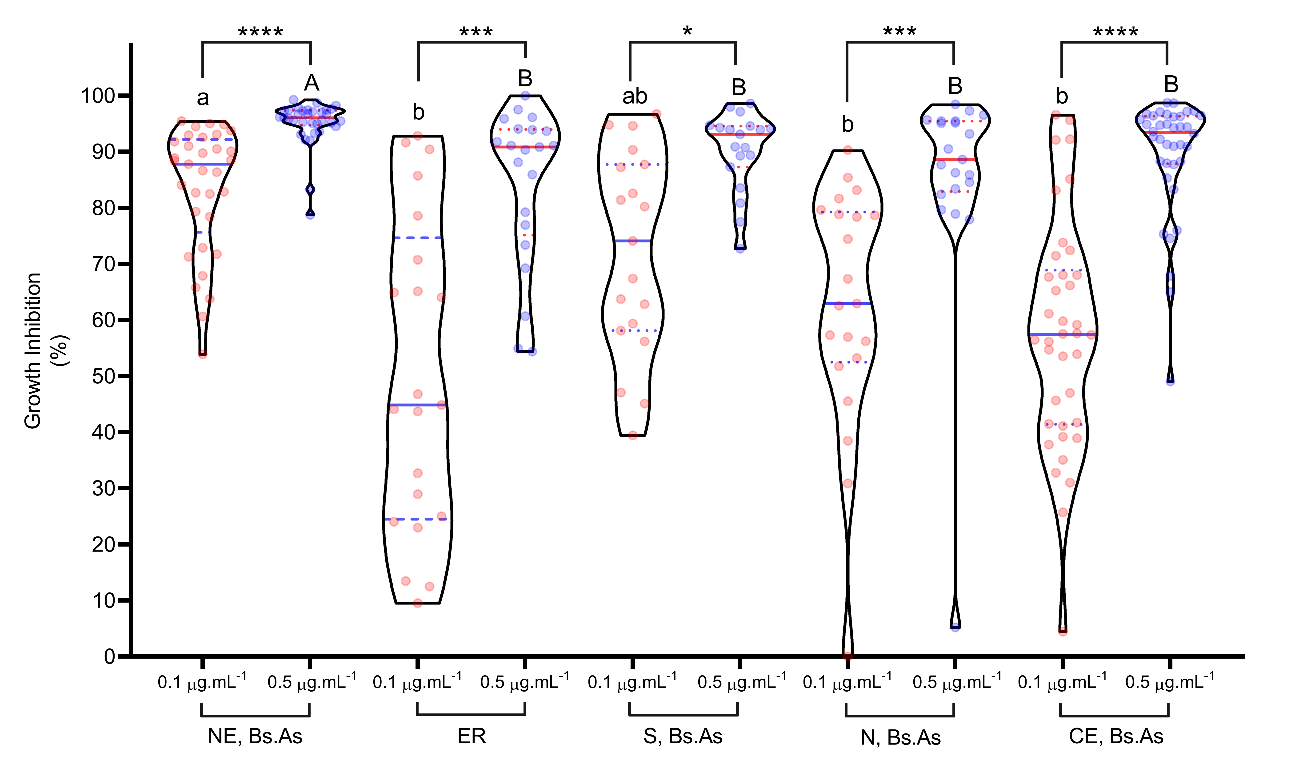

**Supplementary Fig 1**. **Effect of prothioconazole on mycelial growth inhibition in 133 *L. maculans* isolates from five geographic regions of Argentina**. The frequency of mycelial growth inhibition in response to two discriminatory doses (0.1 and 0.5 µg mL^-1^) of prothioconazole in PDA medium is shown. Growth inhibition is expressed as a percentage relative to a control without prothioconazole. The violin plot illustrates the distribution of inhibition levels among isolates for each dose. Normality was assessed using the Shapiro-Wilks test, confirming a non-normal distribution. A non-parametric Mann-Whitney test was applied within each region, revealing significant differences (P ≤ 0.0001), indicated by asterisks. Lowercase letters denote comparisons among regions at 0.1 µg mL^-1^, where different letters indicate significant differences. Uppercase letters represent comparisons at 0.5 µg mL^-1^, with different letters indicating significant differences among regions.
