## Supplemental Figure 2 for "Sensitivity of Argentinean isolates of *Leptosphaeria maculans* to Azoxystrobin, Boscalid and Prothioconazole"

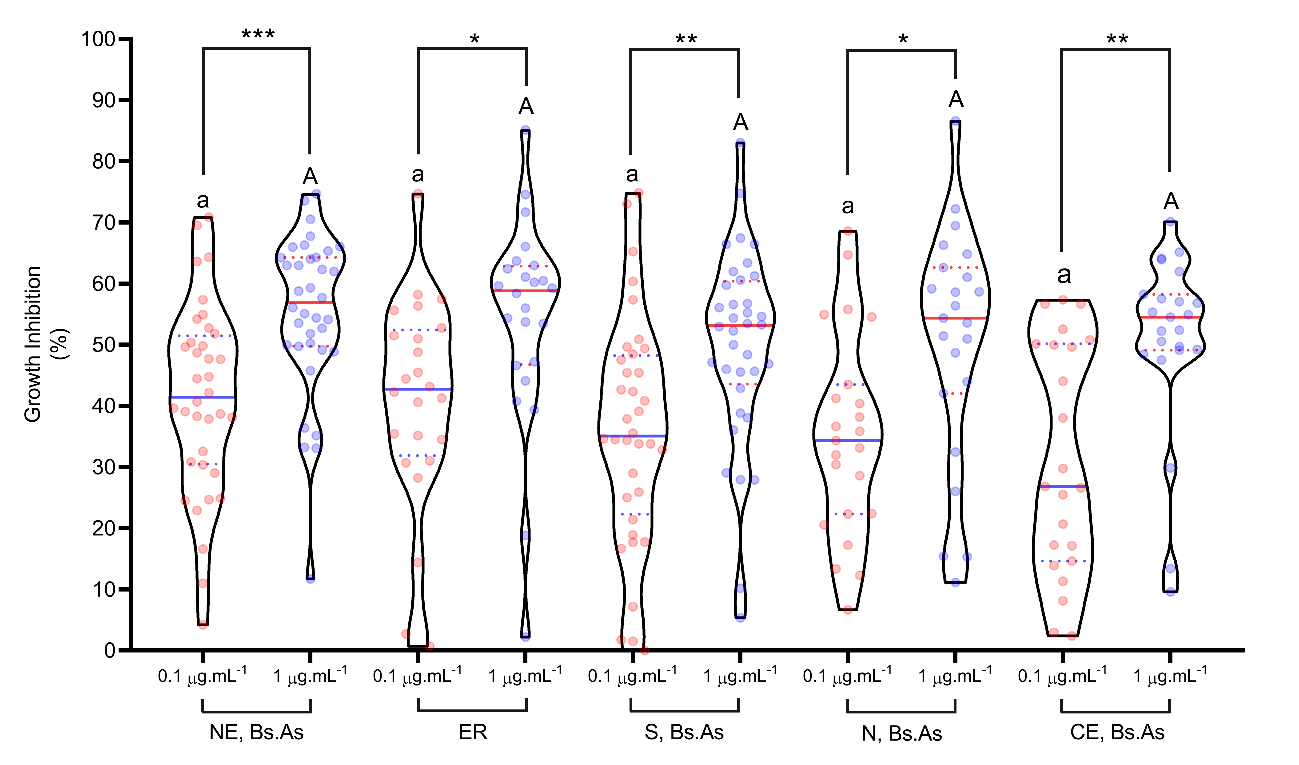

**Supplementary Figure 2**. **Effect of azoxystrobin on mycelial growth inhibition in 142 *L. maculans* isolates from five geographic regions of Argentina**. The inhibition of pycnidiospore germination and mycelial growth, measured photometrically at 405 nm, was assessed using two discriminatory doses (0.1 and 1 µg mL^-1^) of azoxystrobin in liquid V8 medium. Growth inhibition is expressed as a percentage relative to a control without azoxystrobin. The violin plot shows the distribution of inhibition levels among isolates for each dose. The Shapiro-Wilks test confirmed a non-normal distribution, so a non-parametric Mann-Whitney test was performed within each region, revealing significant differences (P ≤ 0.0001), indicated by asterisks. Lowercase letters denote comparisons among regions at 0.1 µg mL^-1^, while uppercase letters indicate comparisons at 1 µg mL^-1^. Different letters represent significant differences between regions.
