## Supplemental Figure 3 for "Sensitivity of Argentinean isolates of *Leptosphaeria maculans* to Azoxystrobin, Boscalid and Prothioconazole"

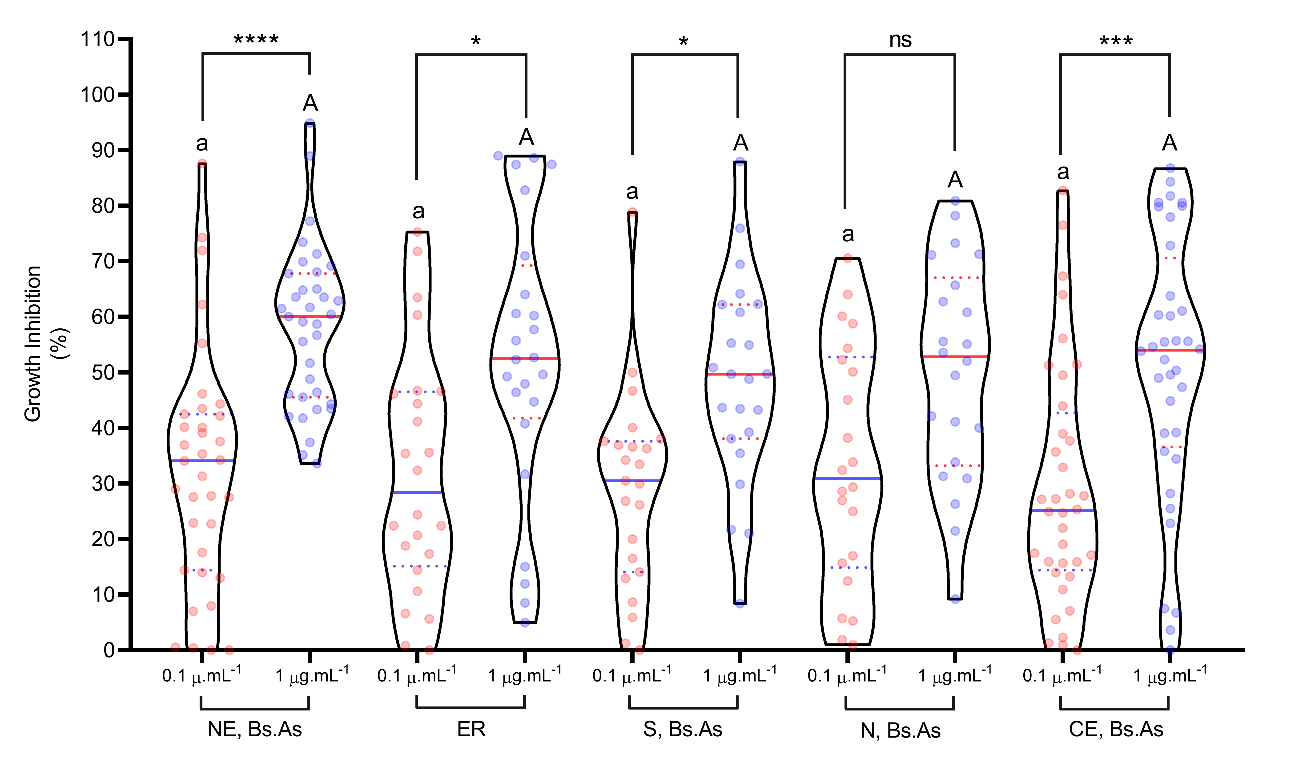

**Supplementary Figure 3.** **Effect of boscalid on mycelial growth inhibition in 140 *L. maculans* isolates from five geographic regions of Argentina**. Pycnidiospore germination and mycelial growth inhibition were measured photometrically at 405 nm in response to two discriminatory doses (0.1 and 1 µg mL-1) of boscalid in liquid V8 medium. Growth inhibition is expressed as a percentage relative to a control without boscalid. The violin plot illustrates the distribution of inhibition levels among isolates for each dose. The Shapiro-Wilks test confirmed a non-normal distribution, so a Mann-Whitney test was performed within each region, revealing significant differences (P ≤ 0.0001), indicated by asterisks. Lowercase letters indicate comparisons among regions at 0.1 µg mL^-1^, while uppercase letters correspond to 1 µg mL^-1^. Different letters represent significant differences between regions.
