## Supplemental Table 1 for "Sensitivity of Argentinean isolates of *Leptosphaeria maculans* to Azoxystrobin, Boscalid and Prothioconazole"

**Supplementary table 1**. Spearman correlation analysis of *L. maculans* isolates with RSI < 1 for at least one fungicide. The table shows correlation coefficients (r) and p-values for sensitivity comparisons at low and high doses. Significant correlations (p < 0.05) are marked with (*).

| **Comparison** | **Number of isolates** | **Spearman *r*** | ***P* - value** | **Significance** |
| --- | --- | --- | --- | --- |
| ***Prothioconazole 0.1 µg mL^-1^ vs Azoxystrobin 0.1 µg mL^-1^*** | 27 | 0.045 | 0.823 | ns |
| ***Prothioconazole 0.5 µg mL^-1^ vs Azoxystrobin 1 µg mL^-1^*** | 13 | -0.093 | 0.732 | ns |
| ***Prothioconazole 0.1 µg mL^-1^ vs Boscalid 0.1 µg mL^-1^*** | 27 | -0.011 | 0.947 | ns |
| ***Prothioconazole 0.5 µg mL^-1^ vs Boscalid 1 µg mL^-1^*** | 14 | -0.032 | 0.882 | ns |
| ***Azoxystrobin 0.1 µg mL^-1^ vs Boscalid 0.1 µg mL^-1^*** | 39 | 0.318 | 0.042 | * |
| ***Azoxystrobin 1 µg mL^-1^ vs Boscalid 1 µg mL^-1^*** | 17 | 0.142 | 0.517 | ns |
